# Mathematical modeling of ion homeostasis & cell volume stabilization: impact of ion transporters, impermeant molecules, & Donnan effect

**DOI:** 10.1101/2022.12.08.519683

**Authors:** Zahra Aminzare, Alan R. Kay

## Abstract

The pump-leak mechanism (PLM) first, described by Tosteson and Hoffman (1960), demonstrates how the activity of the *Na*^+^ − *K*^+^ ATPase (NKA) can counteract the osmotic influx of water stimulated by the presence of impermeant intracellular molecules. We derive analytical solutions for the steady state ion concentrations, voltage, and volume of a cell, by including impermeant extracellular molecules, variable impermeant charge, and Cation-Chloride Co-transporters (CCC). We demonstrate that impermeant extracellular molecules could stabilize a cell without NKA activity but argue that it is unlikely to play a significant role *in vivo*. Significantly we have shown that the precise form of the NKA is unimportant for determining the steady state in PLMs. We have derived an analytical expression for the steady state of the PLM with one of the Cation-Chloride Co-transporters, either KCC, NCC, or NKCC, active. Notably, we have demonstrated that NCC at high pump rates can destabilize cells, which could account for the rarity of this co-transporter. In addition, we show that the reversal of any of the CCCs is unlikely. Importantly, we link the thermodynamics of the NKA to the PLM to show that there is a natural limit to the energy utilized by the PLM that prevents futile cycles. We show that the average charge on the intracellular impermeant molecules influences ion distributions but has no impact on energy utilization. Our study shows that analytical mathematical solutions from physically well-grounded models provide insight into ion transport systems that could only be obtained from numerical simulations with great difficulty.

**Significance Statement:** The regulation of cell volume is fundamental to the stability of all tissue. Animal cells regulate their volume by actively pumping sodium and potassium ions, preventing the water’s osmotic influx from blowing up the cell. Based on the physical laws that determine ion and water fluxes, we derive equations that allow one to predict how pump rates and ion conductances combine to stabilize cell volume. The action of the sodium pump consumes about 30% of a cell’s energy budget, and we demonstrate the rate of ion pumping is constrained so that cells do not consume excessive energy. Our work also demonstrates the power of closed-form mathematical equations in characterizing such pump-leak systems.

## I. Introduction

Because cells require a large inventory of molecules to function, and since they are housed in water permeable membranes, they are confronted with an unrelenting osmotic challenge. The presence of impermeant molecules within a cell establishes a Donnan (or Gibbs-Donnan) effect that, if left unchecked, will lead to the cell volume increase due to the osmotic flux of water until it lyses. The Donnan effect can be counteracted by establishing a cell wall that allows the development of a turgor pressure, which is what plants and bacteria do. Animals evolved a strategy, the pump-leak mechanism (PLM)[1], whereby *Na*^+^ is actively pumped out of cells, while *K*^+^ is pumped in. This effectively stabilizes the cell by equalizing the osmolarity across the membrane but requires continuous energy expenditure to preserve this dynamic steady state. The operation of the PLM leads to the marked asymmetry in *Na*^+^ and *K*^+^ concentrations inside and outside cells and to the development of negative membrane potential.

In 1911, Frederick Donnan showed theoretically how intracellular impermeant molecules could influence ion distributions, water fluxes, and voltage within a cell [2]. The influence of the impermeant molecules within a cell on ion and water distributions and membrane voltage is precisely the Donnan effect. The presence of these molecules served as the spur driving cells to evolve active strategies for counteracting its effect, which, if left unopposed, leads to cell rupture. The effect of the PLM is to effectively hold the Donnan effect at bay, forestalling a Donnan equilibrium.

The ion transport properties of a cell are typically modeled by measuring or abstracting from the literature the characteristics of the ion channels and transporters, incorporating them into a model, and determining if the model recapitulates the physiology of the cell. In contrast, we are attempting to assess the global characteristics of a set of ion channels and transporters, not just single instances. In our case *Na*^+^, *K*^+^, and *Cl*^−^ ion channels, the *Na*^+^*/K*^+^ − ATPase (NKA) and three kinds of SLC12 transporters are considered. It is an attempt to view whether qualitatively different behaviors exist in the model’s parameter space. Conventional simulations can recapitulate behavior; our approach aims to provide insight into the operation of such cells.

For the cell systems that we consider here, there are only three possible solutions: (1) The cell volume increases monotonically until the cell bursts. What happens mathematically is that the volume tends to infinity. (2) The cell collapses to zero volume. (3) The cell volume stabilizes at a constant finite volume. We will refer to this as the **steady state** solution, which is characterized by the concentrations of all molecules within the cell, the transmembrane voltage (*V*), and the cell volume (*w*).

Cation-chloride co-transporters (CCCs) are a family (SLC12) of the Solute Carrier transporter superfamily, which transport *Na*^+^ and/or *K*^+^ and *Cl*^−^ across the cell membrane [3]. There are three types of electroneutral CCCs: the *Na*^+^– *Cl*^−^ co-transporters, NCC; the *Na*^+^– *K*^+^– *Cl*^−^ co-transporters, NKCC; and the *K*^+^– *Cl*^−^ co-transporters, KCC. The CCCs are secondary transporters that depend on the nonequilibrium ion gradients established by the NKA to drive *Cl*^−^ out of equilibrium and perturb the equilibria of *Na*^+^ and *K*^+^. Hence, in the absence of an active NKA, the CCCs cannot drive ion concentrations out of equilibrium. The SLC12 family is involved in a large number of aspects of physiology, and the literature is vast, so we will not attempt to summarize it, but we point the reader to recent summaries [4], [5].

An important part of modeling the transport characteristics of cells is incorporating water permeability. Doing this requires that one include the mechanics of the cell since differences in osmolarity across the plasma membrane will allow water fluxes and hence volume changes. Moreover, once one introduces water fluxes, it opens the possibility that the cell might not be stable since they can now shrink or lyse [6]. Most ion transport models, with some notable exceptions, do not include water fluxes, which would mask any instabilities that might arise. Part of the objective of this study is to call attention to the importance of including the osmotic flux of water in any models that attempt to model ionic homeostasis in cells.

The pump-leak equations (PLEs) are a set of four differential equations and one algebraic equation that typically model the PLM. These equations describe the dynamics on *Na*^+^, *K*^+^, and *Cl*^−^, the cell volume, and the membrane potential (see Section II for more details). PLEs were first developed in [1] and have been employed and extended by others to understand cell volume control in single cells [6], [7], [8], [9], [10] and epithelial [11], [12], [13]. In [14], Keener and Sneyd derived a very useful analytic solution to the PLEs. In [15], Mori established analytical results on the existence and stability of steady states for a general class of PLEs. Here, we have extended PLEs to include impermeant extracellular molecules and Cation-Chloride Co-transporters. We show that the former can help establish an equilibrium that requires no energy to sustain. From this passive case, we then move to the active case with an NKA.

There is a range of models for the NKA, from a simple constant model to complex nonlinear models [14], [16], [17], [18]. We will show the form of these models does not affect the steady-state value of a cell, and therefore, considering a constant rate model for the NKA pump suffices. Operation of the NKA requires ATP, and estimates are that it expends something like 20-30% of a cell’s energy [19]. Using our equations, we show how cellular conductances and pump stoichiometry influence energy expenditure to preserve a constant volume. In the last third of the paper, we examine the effect of the CCCs on ion distributions and cellular stability. Interestingly, we show that the NCC has the capability of destabilizing cells, which might explain its restricted distribution.

The rest of the paper is organized as follows: In Section II, we introduce the PLEs for a single cell. In the following sections, we examine the effect of three different mechanisms on cell stabilization. In Section III, we consider a passive cell (i.e., one where the NKA is not active) and show how extracellular impermeant molecules stabilize a cell. In Section IV, we show how an NKA pump can regulate cellular stability. In Section V, we incorporate three co-transporters into the model of an active cell and explore their effect on cell volume regulation. Finally, we conclude in Section VI and provide more details in Sections VII and VIII.

## II. Cell model

In this paper, we examine the conditions that will stabilize the volume of a cell or work against it, using the Pump-leak mechanism (PLM). As shown in Figure 1 we consider a simple cell with a pliant membrane that is permeable to *Na*^+^, *K*^+^, *Cl*^−^, and water, immersed in an extracellular medium that has a fixed concentration. We will assume that there are no concentration differences within the cell since diffusion is rapid within a small cell. In this work, we neglect pH, and hence 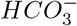 for simplicity; we will include them in future explorations and note the work of Li *et al*. [20] in this regard. We further assume that concentrations and voltages are uniform, showing no spatial gradients. For most single cells, this is a realistic assumption since diffusion is rapid on the micron scale. For two adjacent cells coupled through the cleft gap, spatial variability becomes essential [21], [22].

**Fig. 1:**
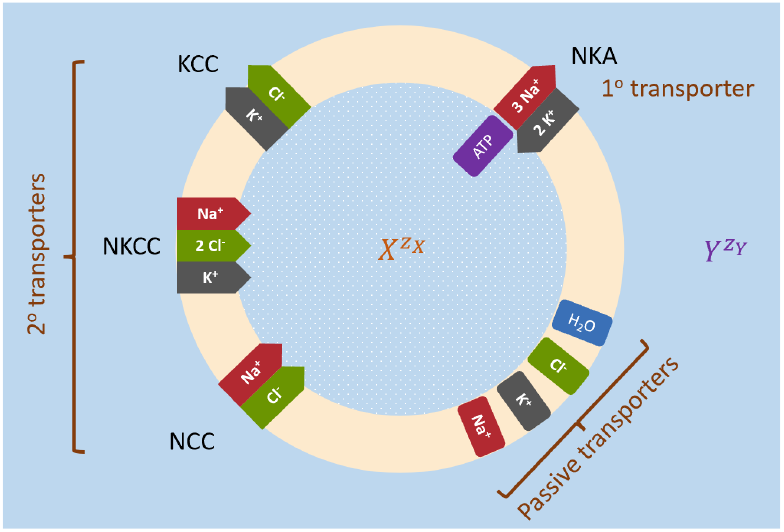
Schematic of the model cell with *Na*^+^, *K*^+^, *Cl*^−^ and water channels, NKA pump (primary transporter), KCC, NCC, and NKCC (secondary transporters), and intracellular (*X*) and extracellular (*Y*) impermeant molecules.

To begin, we review the construction of the PLM and show its firm grounding in physical principles. There are two major physico-chemical forces that are at play across the membrane and exert a significant role in determining cell volume. First, water moves across the plasma membrane if there is a gradient of osmolarity across the membrane. Second, the interaction between charges in the solution, governed by Maxwell’s laws, ensures that no significant imbalances in charge arise [23]. We write down a set of equations that determine how ions move in and out of the cell.

The PLEs are derived from physical laws, which in conjunction with the system parameters, determine the dynamics of the system. Mori [15] proved that the PLEs with a constant pump obey a free-energy relationship or extremum principle. That is, for some values of its parameters, the system can be represented by an energy surface, with a unique minimum corresponding to the steady state. For other values of the parameters, the cell either bursts or shrinks to zero volume, but no oscillations (i.e., limit cycles) are possible. For all possible steady states, the osmotic pressure inside equals that outside, and the intracellular solution is approximately electroneutral. These are sometimes referred to as constraints, but they are simply the outcome of the energetics of the system. Using the notion of constraint makes it seem like the system behaves teleologically.

### A. The Pump-Leak Equations (PLEs)

Since the membrane is permeable to water, if there are differences in osmolarity across it, water will move by osmosis. The osmolarities of the intra- and extracellular solutions are respectively 𝒪_*i*_ and 𝒪_*e*_:

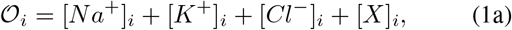

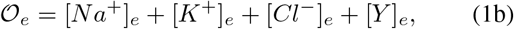

where [ion]_*i*_ and [ion]_*e*_ represent the intracellular and extra-cellular concentrations, respectively. [*X*]_*i*_ represents all impermeant intracellular molecules, which include metabolites and macromolecules. Similarly, [*Y*]_*e*_ represents the extracellular impermeant molecules. Because osmosis is a colligative effect, one need only consider the number of moles of these molecules. Note that we assume [*Y*]_*e*_ is constant and the number of moles of the intracellular impermeant molecule is constant, i.e., *X*_*i*_ is constant. For the rest of the paper, we assume that *X*_*i*_ > 0. Otherwise, when *X*_*i*_ = 0, the cell volume, *w*, shrinks, i.e., *w* →0.

The osmotic flux of water, and hence the change in volume, denoted by *w*, is governed by Starling’s equation:

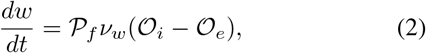

where *P*_*f*_ is the osmotic permeability and *ν*_*w*_ is the partial molar volume of water.

The total concentration of charge inside and outside the cell is given by:

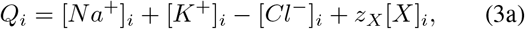

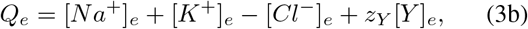

where *z*_*X*_ and *z*_*Y*_ are the average charge of *X*_*i*_ and *Y*_*e*_, respectively.

Because of the energetic cost of separating charges, isolated solutions will have a net charge of zero, so we can set *Q*_*e*_ = 0. The membrane potential of the cell, denoted by *V*, can be modeled exactly by the following algebraic equation [24]:

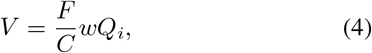

where *F* is Faraday’s constant and *C* is the total cell capacitance. *C* = *C*_*m*_ × *A*_0_ where *C*_*m*_ is the unit membrane capacitance (≈ 1*μFcm*^−2^) and *A*_0_ is the cell surface area.

Note that by (4),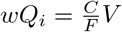. Since for small *C*, the ratio 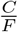 is close to zero, if *w* is finite, then *Q*_*i*_ ≈ 0. Therefore, once we assume *Q*_*e*_ = 0, we can conclude that approximately

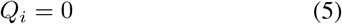

while *V* does not need to be zero.

The rates at which a particular ion flows into or out of the cell are determined by the net flux through its ion channels (Ohm’s law) and active transporter(s). The following three differential equations describe the dynamics of three main ion concentrations in a cell, namely *Na*^+^, *K*^+^, and *Cl*^−^:

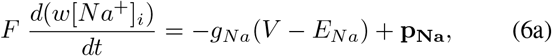

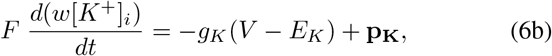

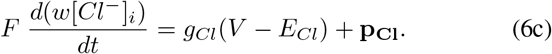

In these equations, *Eion* is the Nernst potential of an ion and is equal to

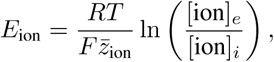

where *RT* is the ideal gas constant times absolute temperature, 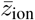 is the corresponding valence, and *g*_ion_ is the corresponding ionic conductance. We assume that the total number of ion channels on the cell membrane remains unchanged. Hence, the total conductances stay constant. **p**_**Na**_, **p**_**K**_, and **p**_**Cl**_, represent the current generated by ion transporters. The precise nature of these components will be taken up in the following sections.

We will call (2), (4), and (6) the pump-leak equations (PLEs) [14]. In [15], [25], the authors studied the PLEs and employed the NKA pump as a mechanism to stabilize the volume of a single cell. Here, we generalized the PLEs by incorporating the Cation-Chloride co-transporters and impermeant molecules in extracellular fluid. Our goal is to understand the effect of each of these mechanisms on controlling the stability of a single cell.

All the parameter values are given in Table I unless otherwise stated.

**TABLE I.** Parameters used in this paper

| Params./Var. | Description | Value |
| --- | --- | --- |
| $R_0$ | initial radius | $5 \mu m$ |
| $A_0$ | initial area | $4\pi R_0^2$ |
| $W_0$ | initial volume | $\frac{4}{3}\pi R_0^3$ |
| $R$ | Universal gas const. | $8.31 \text{ (C/V)/(mol K)}$ |
| $T$ | Temperature | $310 \text{ K (37 C)}$ |
| $F$ | Faraday const. | $96485 \text{ C/mol}$ |
| $C$ | Total capacitance | $0.0001 A_0 \text{ (F)}$ |
| $\mathcal{P}_f \nu_w$ | Osmotic permeability<br>× partial molar volume | $0.0126 A_0$ |
| $g_{Na}$ | Sodium conduct. | $0.001 A_0 (S)$ |
| $g_K$ | Potassium conduct. | $0.03 A_0 (S)$ |
| $g_{Cl}$ | Chloride conduct. | $0.002 A_0 (S)$ |
| $\mathcal{O}_e$ | Extra. osmolarity | $300 \text{ mM}$ |
| $[Y]_e$ | Extra. impermeant molecule | $1 \text{ mM}$ |
| $z_Y$ | Extra. valence | $-1$ |
| $[K^+]_e$ | Extra. Potassium concent. | $3 \text{ mM}$ |
| $[Cl^-]_e$ | Extra. Chloride concent. | $149 \text{ mM}$ |
| $[Na^+]_e$ | Extra. Sodium concent. | $147 \text{ mM}$ |
| $X_i$ | Intra. impermeant molecule | $50 \times W_0 \text{ mM} \times \text{dm}^3$ |
| $z_X$ | Intra. valence | $-1$ |
| $\nu$ | sodium stoichiometry | $3$ |
| $\kappa$ | potassium stoichiometry | $2$ |
| $p_{NKA}$ | NKA pump const. | $p \times A_0$ |
| $p_{KCC}$ | KCC co-transporter const. | $c_{KCC} \times A_0$ |
| $p_{NCC}$ | NCC co-transporter const. | $c_{NCC} \times A_0$ |
| $p_{NKCC}$ | NKCC co-transporter const. | $c_{NKCC} \times A_0$ |

## III. Passive cells: The effect of extracellular impermeant molecules on cell stabilization

In this section, we consider a passive cell ((2), (4), and (6) without any pumps or co-transporters) and analytically study the effect of extracellular impermeant molecules on cellular stability. To this end, we compute the equilibrium values and explore the effect of [*Y*]_*e*_ and *z*_*Y*_ on them.

To keep the extracellular osmolarity 𝒪_*e*_ constant and ensure the electroneutrality, we assume that [*K*^+^]_*e*_ is fixed, and [*Na*^+^]_*e*_ and [*Cl*^−^]_*e*_ vary as follows:

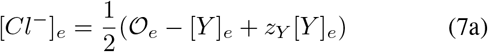

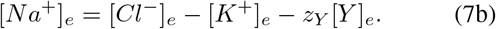

We will show that for a passive cell, namely one that does not expend energy, a stable thermodynamic equilibrium can be attained simply by the provision of impermeant extracellular molecules ([*Y*]_*e*_ > 0), which stabilizes the volume of the cell.

To find the equilibrium, we set the derivatives in (2) and (6) to zero and consider (4) as a constraint. Setting the derivatives in (6) to zero gives:

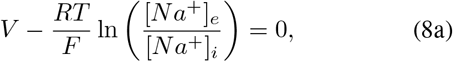

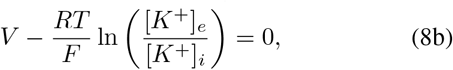

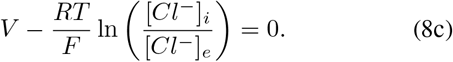

Using (8), we have:

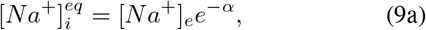

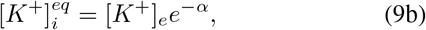

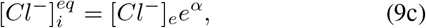

where 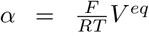 and the superscript *eq* stands for equilibrium. This is a stable Donnan equilibrium.

A further condition for equilibrium is that there be no change in the volume, that is, the derivative in (2) is zero. Therefore the intracellular osmolarity becomes equal to the extracellular osmolarity:

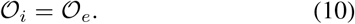

In what follows, using the Donnan equilibrium, osmotic equilibrium, and electroneutrality assumption, we derive an expression for 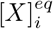 in terms of *z*_*Y*_, [*Y*]_*e*_, and *O*_*e*_. Then, we use 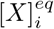 and the fact that 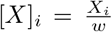, and derive an expression for *w*^*eq*^ in terms of the extracellular quantities.We write [*X*]_*i*_ and *w* in terms of the extracellular quantities because the extracellular quantities are assumed to be fixed.

Following Fraser and Huang [8], the equilibrium of intracellular concentrations can be described as the extracellular concentrations and *z*_*X*_ as follows.

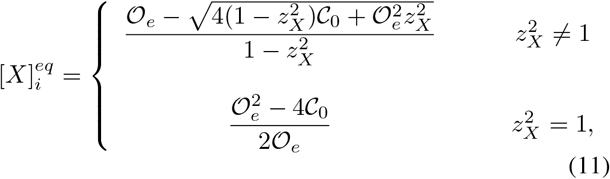

where 𝒞_0_ = [*Na*^+^]_*e*_[*Cl*^−^]_*e*_ + [*K*^+^]_*e*_[*Cl*^−^]_*e*_. The equilibrium values of the intracellular ion concentrations, cell volume, and voltage are equal to

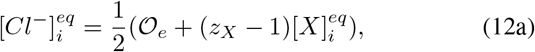

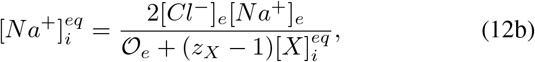

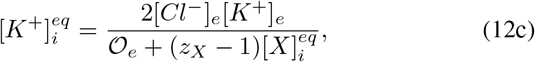

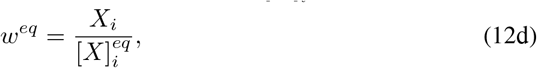

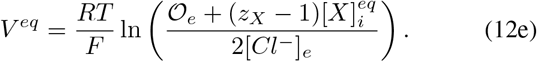

In what follows, we show that [*Y*]_*e*_ can be used as a mechanism to stabilize the volume of a passive cell. To this end, we rewrite (11) as a function of [*Y*]_*e*_.

**Case 1**. Assume *z*_*X*_ = ± 1. Then,

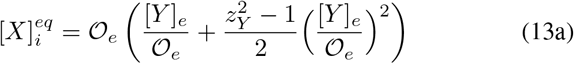

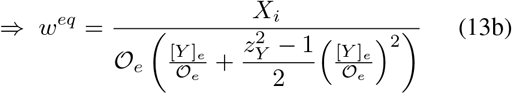

**Case 2**. Assume *z*_*X*_ ±1. Then, 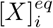 becomes

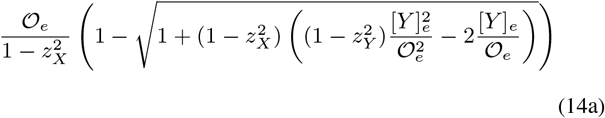

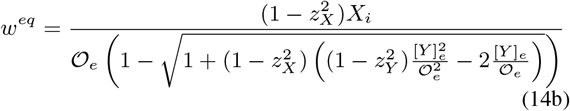

Substituting [*Cl*^−^]_*e*_, [*Na*^+^]_*e*_ (from (7)) and 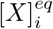 (from (13) or (14)) into (12), we obtain new expressions for the equilibrium values of the ion concentrations and voltage in terms of *z*_*Y*_, [*Y*]_*e*_, and fixed parameters [*K*^+^]_*e*_ and 𝒪_*e*_. We do not show the new formulae here but plot them in Figure 2.

**Fig. 2:**
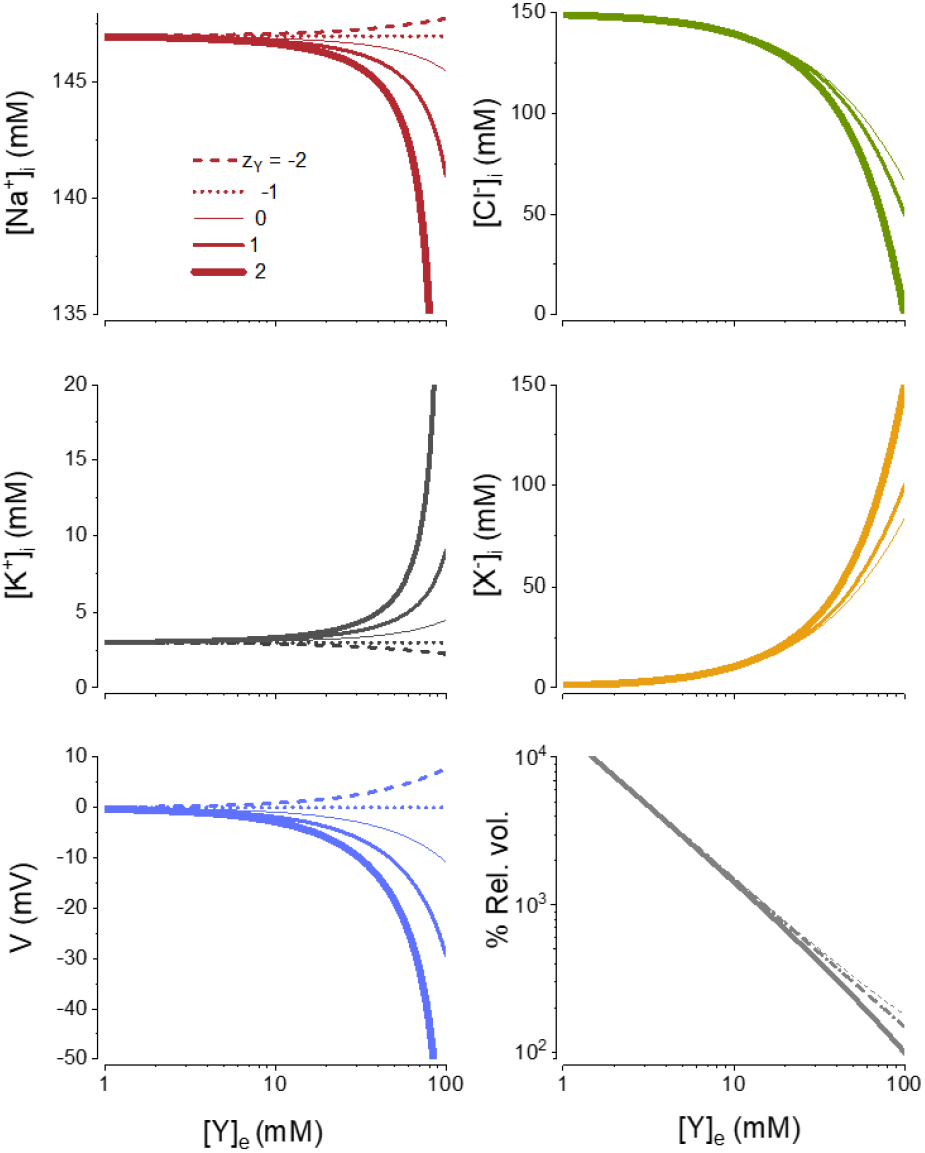
The effect of *Y*_*e*_ on the steady state of a passive cell. The equilibrium values of ion and intracellular impermeant molecule concentrations, volume, and voltage are plotted as functions of [*Y*]_*e*_ for 5 different values of *z*_*Y*_ : −2, −1, 0, 1, 2. These equilibria are calculated from (12) and (13). Volumes are normalized by the minimum steady state volume of the cell for *Y*_*e*_ = 2.

Reciprocal dependence of *w*^*eq*^ on 𝒪_*e*_ and [*Y*]_*e*_/ 𝒪_*e*_ implies that for a constant 𝒪_*e*_, as [*Y*]_*e*_ increases, *w*^*eq*^ decreases. To see this effect, in Figure 2, we plot the equilibrium values of ion concentrations, impermeant molecule, volume, and voltage as [*Y*]_*e*_ changes. Each equilibrium is plotted for 5 different values of *z*_*Y*_ : −2, −1, 0, 1, 2. In this figure, *z*_*X*_ = −1. Qualitatively, the behavior of the cell is similar for *z*_*X*_ = 1 or *z*_*X*_ ≠ ±1 (the simulations are not shown here). Also, when [*Y*]_*e*_ = 0, *w*^*eq*^ = ∞. Therefore, to obtain a finite equilibrium *w*^*eq*^, we assume that [*Y*]_*e*_ ≠ 0 and use it as a mechanism to stabilize a cell in the absence of any pumps. Note that a small amount of [*Y*]_*e*_ (e.g., [*Y*]_*e*_ = 1 mM, 1/3 of extracellular potassium concentration) tremendously reduces the cell volume. See Figure 2.

Based on (13) and Figure 2, the chloride concentration and volume always decrease as *Y*_*e*_ increases. For *z*_*Y*_ *<* −1, sodium and voltage are increasing functions of *Y*_*e*_ while potassium is a decreasing function of *Y*_*e*_. For *z*_*Y*_ = −1, sodium, potassium, and voltage are constant functions of *Y*_*e*_. For *z*_*Y*_ > −1, sodium and voltage become decreasing functions of *Y*_*e*_ while potassium becomes an increasing function of *Y*_*e*_. Since, in the absence of an NKA pump and extracellular impermeant molecules, a cell bursts, for the rest of the paper, we fix [*Y*]_*e*_ = 1 mM and *z*_*Y*_ = −1.

## IV. Active cells: The effect ofnka on cell stabilization

In this section, we include the active transport of *Na*^+^ and *K*^+^ by incorporating a model of the NKA into the cell model, (6). The stoichiometry of the NKA is 3*Na*^+^ : 2*K*^+^. In this paper, we will consider alternative stoichiometries. We assume that for each molecule of ATP hydrolyzed, the pump transports *νNa*^+^ ions out of the cell and *κK*^+^ ions into the cell. Then, by (6),

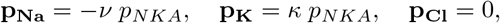

where *p*_*NKA*_ = *pA*_0_ and *p* is the density of NKA multiplied by the rate of ATP hydrolysis by a single pump. A great number of models have been put forward to capture the kinetics of the NKA. However, as we show in Appendix VIII-A, the precise form of the pump model does not have an impact on the steady state behavior of the PLM. Hence, we use the simplest form of a model, a constant value for *p*, which we refer to as the **constant** NKA model (in the sense that it is independent of our model variables such as ion concentrations, etc.), although we vary its magnitude to explore its impact on the steady state behavior of the cell. In what follows, we compute the steady state values of the PLM analytically and explore the effect of various model parameters on these steady states.

Keener and Sneyd [14] derived equations for the steady state for the case of *z*_*X*_ ≤ −1. Later, Mori [15] relaxed this condition and proved the existence of a steady state for any *z*_*X*_. Here, we extend their work by including impermeant extracellular molecules, arbitrary *z*_*X*_, and arbitrary NKA stoichiometry. We show that the steady states exist for a specific range of *p*_*NKA*_; see (18) below.

### A. Steady states of NKA-pump leak model

Similar to Section III, we compute the steady state values of PLM, which include the concentrations 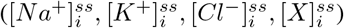, volume (*w*^*ss*^) and voltage (*V* ^*ss*^). Since the cell is active now, we replace the superscript *eq* with *ss*, which stands for steady states. The steady state of the cell is given by the following equations:

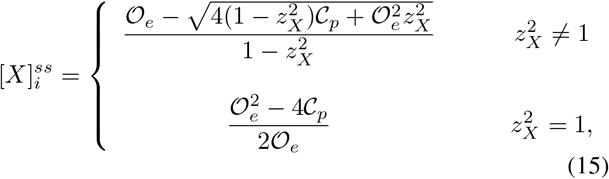

where

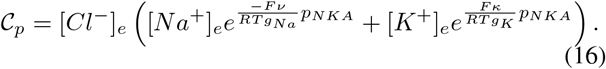

The intracellular ion concentrations, cell volume, and voltage at the steady state are equal to

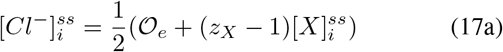

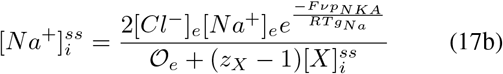

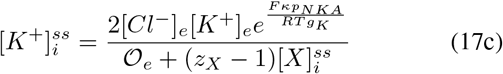

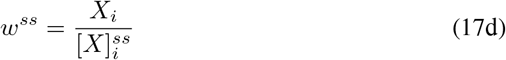

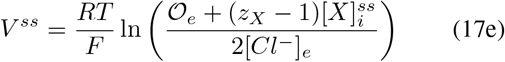

where 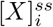 is as given in (15). When *p*_*NKA*_ = 0, the above equations become identical to the ones for a passive cell, as given in (11) and (12).

To establish a cell with a stable finite volume, 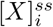 must be positive. To have a positive 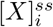, the NKA-pump rate *p*_*NKA*_ = *pA*_0_ must satisfy the following condition

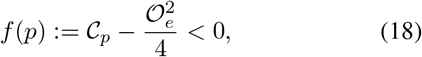

where 𝒞_*p*_ is defined by (16). It is clear that *f* (0) < 0 and the graph of *f* is first decreasing and then becomes monotonically increasing. Therefore, there is a *p*_*NKA*_ = *p*_max_ at which the function *f* changes sign from negative to positive. For the parameters presented in this paper *p*_max_ ≈ 0.00157 *A dm*^−2^. Note that the condition given in (18) is independent of *z*_*X*_.

In summary, we have shown that for *p* < *p*_max_/*A*_0_, there exists a unique steady state given in (15)–(17e). Without computing the steady states, in [15, Propositions 3.3 & 3.4], Mori proved that PLEs with a constant pump that satisfies (18) (which is equivalent to [15, Eq. (3.5)]) admits a unique and globally asymptotically stable steady state. This means that if any of the variables are perturbed, the system will return to the steady state. Hence, Mori’s results ensure that the steady state is given in (15)–(17e) is globally asymptotically stable.

### B. Parameters affecting the steady state

Besides the extracellular ion concentrations and osmolarity, which we assume are constant (see Table I), there are four groups of parameters that affect the steady states: (i) the NKA rate; (ii) *z*_*X*_ ; (iii) the NKA stoichiometry; and (iv) the sodium (*g*_*Na*_) and potassium (*g*_*K*_) conductances. In what follows, we discuss the effect of each of these groups of parameters.

#### The effect of NKA rate on the steady state

First, we focus on how the steady state depends upon the pump rate and *z*_*X*_. To do this, we have calculated how the steady state varies as the pump rate is increased. We have previously referred to these as *Cp* plots [25]. In addition, we derive conditions that allow for a stable steady state. We then move on to consider the influence of the pump rate and how a cell could establish an energetically efficient pump rate.

We fix all the parameters at the values given in Table I and vary the NKA rate *p*_*NKA*_ = *pA*_0_. Since *A*_0_ is a constant, we only vary *p*, which is the density of NKA pumps in the membrane, multiplied by the ATP hydrolysis rate of a single pump.

In Figure 3 we plot all the steady state values as functions of *p* where *p* changes from 0 to *p*_max_/*A*_0_. In these calculations [*Y*]_*e*_ = 1 mM, ensuring that in the absence of NKA activity the steady state volume is finite. As is evident from the figure, as the pump rate increases, the steady state volume initially decreases and then increases. This “switchback” was shown in [14] but does not seem to have been remarked on by others. Since it is also observed in the case of nonlinear pump mechanisms (Appendix VIII-A) it appears to be an inherent characteristic of the PLM.

**Fig. 3:**
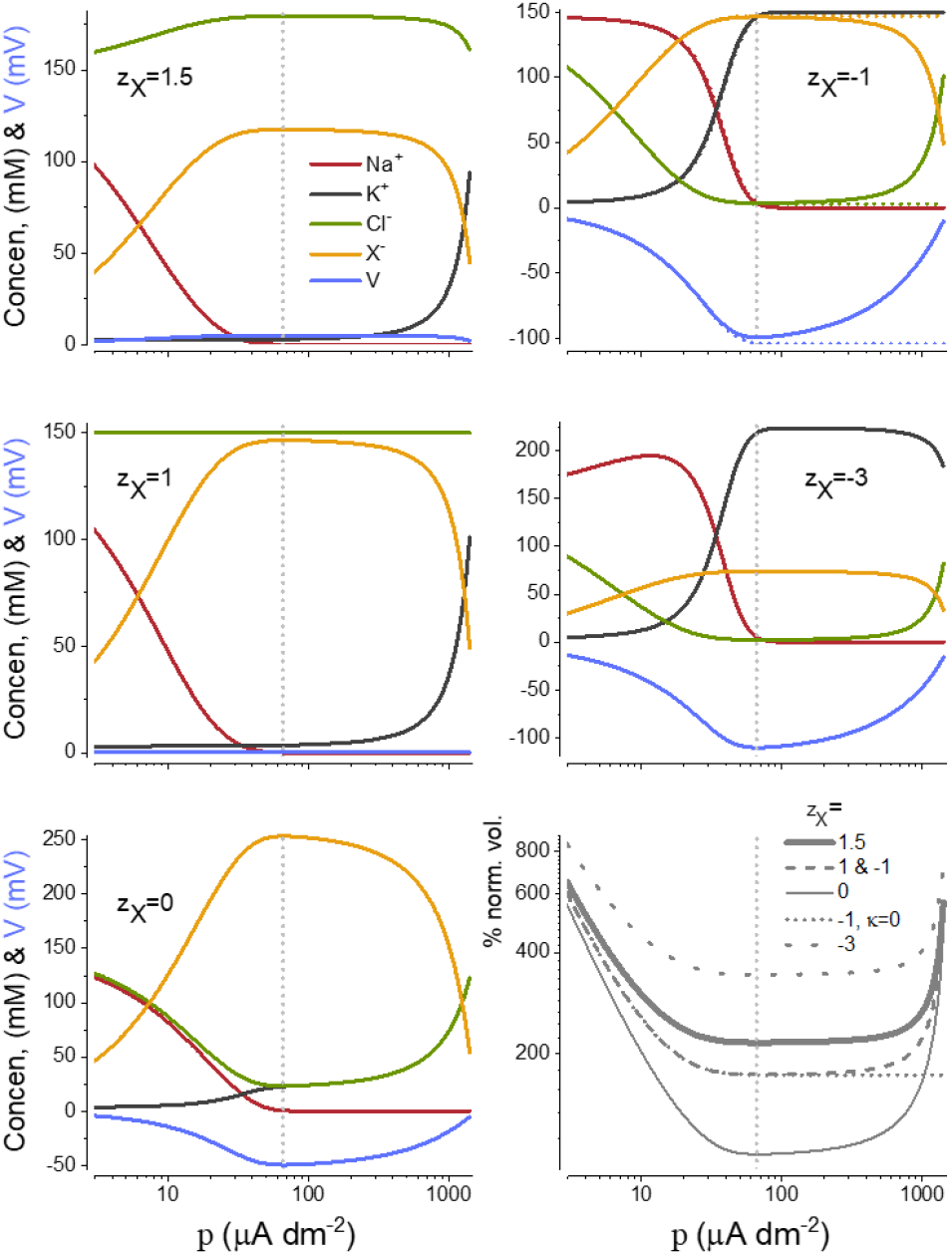
The influence of *z*_*X*_ on the steady state of the PLEs, which are calculated from (17). The steady state values of the variables are plotted as functions of the pump rate at different values of *z*_*X*_ = −3, −1, 0, 1, 1.5. Vertical dotted lines indicate *p* = *p*_*min*_. On the top and bottom right panels (*z*_*X*_ = −1), the concentrations, voltage, and volume are plotted for *κ* = 0 (see the dotted curves). Volumes are normalized by the minimum volume at *z*_*X*_ = 0

It seems likely that cells expend only as much energy as necessary to stabilize cell volume. As the pump rate increases, driving *K*^+^ up and *Na*^+^ down, there comes a point where further increases in pump rate do not change the volume further. In this regime fluctuations in *p* do not lead to large fluctuations in volume, whereas for lower values of *p* they do. For the case where *κ* > 0 there is a minimum inflection point in the *Cp* curves for the volume and voltage. We will call this the *p*_*min*_, which serves as a useful proxy for the optimal pump rate.

To find the value of *p*_min_, we take the derivative of *w*^*ss*^ with respect to *p* and set it equal to zero. Then, for any *z*_*X*_,

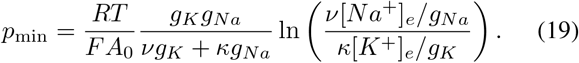

For the cell to have a finite volume, the following condition must hold (as derived in [15, Eq. (4.19)]):

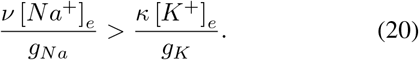

Note that for *κ* = 0, *p*_min_ does not exist. Indeed, when *κ* = 0, the steady state volume is a monotonically decreasing function of *p* (see (17d)). Moreover, when *κ* = 0, [*K*^+^]_*i*_ and [*Cl*^−^]_*i*_ exhibit a Donnan equilibrium for *K*^+^ and *Cl*^−^.

#### The effect of z_X_ on the steady state

The average intracellular valence of impermeant ions, *z*_*X*_, is an important part of the PLM and exerts a strong effect on how ions are distributed across the membrane in the steady state. *z*_*X*_ can be estimated by measuring the concentrations of the predominant permeable cations and anions in the cell; a nontrivial task. Estimates of *z*_*X*_ in several organisms put it in the range of −1.5 to −0.5 [26]. For the rest of the paper, we will choose *z*_*X*_ = −1. *z*_*X*_ is as important as any other physiological measure, like osmolarity, however, it has not attracted much attention and it is unclear whether cells actively attempt to regulate its magnitude.

In Figure 3 we have plotted the *Cp* curves of the PLEs at five different values of *z*_*X*_ (− 3, −1, 0, 1, 1.5). Straight-forward calculations show that 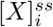 is differentiable with respect to 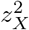 and its derivative is strictly negative. Therefore, as 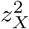 increases 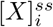 decreases. Note that the sign of *z*_*X*_ doesn’t affect 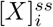. Similarly, the cell volume also depends on 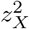 but unlike 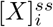, the cell volume is a monotonically increasing function of 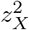.

From (17), we can conclude that the steady state values of [*Cl*^−^]_*i*_ and voltage are decreasing functions of *z*_*X*_ while the steady state values of [*Na*^+^]_*i*_ and [*K*^+^]_*i*_ are increasing functions of *z*_*X*_.

#### The effect of pump stoichiometry on energy consumption & p_min_

There is a direct relationship between the pump rate and ATP consumption; for each cycle of the pump a single ATP is hydrolyzed and hence the energy utilized. In (19), we saw that *p*_*min*_ depends on the stoichiometry of the pump, i.e. the number of *Na*^+^ (*ν*) and *K*^+^ (*κ*) transported per cycle. Since energy is at a premium, there is strong evolutionary pressure on organisms to minimize energy consumption and the pump-leak equations can provide some insight into the factors that influence energy expenditure.

Not all stoichiometries are consistent with a stable cell. For example, if *ν* = 0 and *κ >* 0, from (19), the cell is unstable, as it has been pointed out in [6]. From (19) for a given positive *ν, p*_min_ decreases as *κ* increases, however, there is an energetic limit on how large *κ* can be, which we take up in the next paragraph.

To understand how the stoichiometry of the pump influences the energy of the NKA we will consider the thermodynamics of the pump. To drive a single cycle of the NKA, the hydrolysis of one molecule of ATP is required:

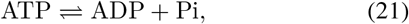

where *Pi* is inorganic phosphate and ADP is adenosine diphosphate. The energy available from this reaction is given by Gibb’s free energy of the reaction [27]:

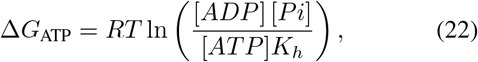

where *K*_*h*_ is the equilibrium constant for the ATP hydrolysis reaction based on a standard state of 1 M. Δ*G*_ATP_ ranges from −50 to −70 kJ mole^−1^ [28]. Following [29], we use the value of Δ*G*_ATP_ = −55 kJ mole^−1^.

The energy required to drive a single cycle of the pump can be calculated from thermodynamics using the chemical potentials of all the species involved [27].

The transport reaction of the pump is:

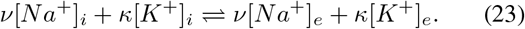

For any ion, the standard formula can be used to calculate its chemical potential. For example, the chemical potential of [*K*^+^]_*i*_ is:

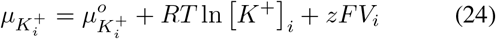

where 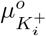 is the standard chemical potential of *K*^+^, and *V*_*i*_ is the intracellular potential.

The Δ*G*_NKA_ for the forward reaction is then:

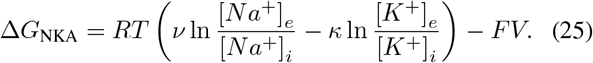

For the pump to function, the following inequality must hold:

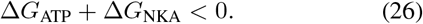

It is worth noting that the pump can only be driven by the hydrolysis of a single ATP, not from the accumulation of energy from the hydrolysis of multiple ATPs. Hence, (26) limits the possible stoichiometries for the pump.

From the free energy relationship given in (25), we can estimate the maximum pump rate *p*_*ATP*_ that is consistent with the energy available from ATP. We can calculate the steady state [*Na*^+^]_*i*_, [*K*^+^]_*i*_, and *V* from the PLEs and use them to compute Δ*G*_NKA_ and determine the value of *p* at which Δ*G*_NKA_ is equal to −Δ*G*_ATP_. The values of *p*_*ATP*_ are shown in Figure 4. If we imagine the NKA pump rate being ramped up, once it reaches *p*_*ATP*_, it can no longer pump. This provides a natural limit that prevents excessive energy consumption.

**Fig. 4:**
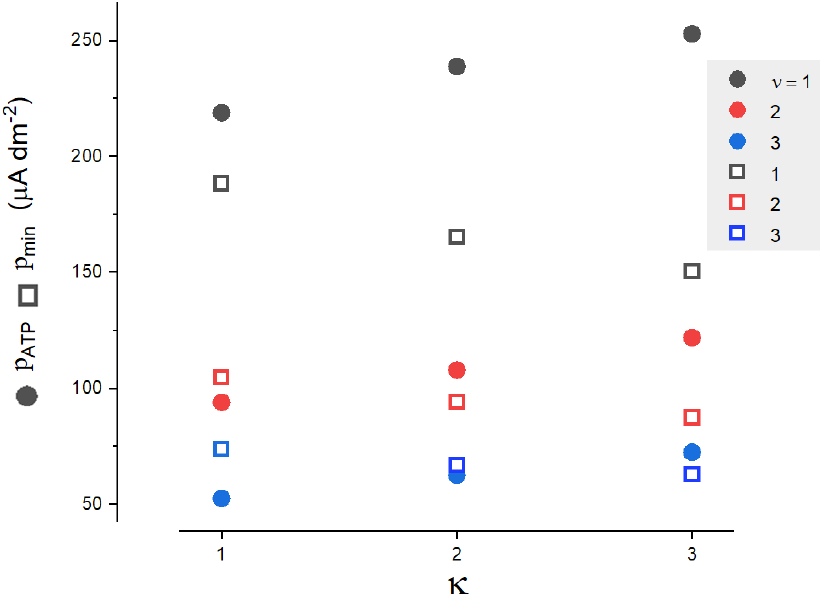
The dependence of *p*_*ATP*_ (solid symbols) and *p*_*min*_ (open symbols) on the stoichiometry of the NKA.

Inserting the steady state values of *Na*^+^ and *K*^+^ from (17) into Δ*G*_NKA_, we can show that for *ν* −*κ* −1 = 0, (26) holds only for *p*_*NKA*_ satisfying

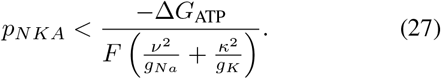

For the given parameters in Table I and for stoichiometry 3*Na*^+^ : 2*K*^+^, (26) holds if *p <* 6.24 *×* 10^−5^, which is of order *p*_max_*/A*_0_ ≈ 17.5 *×* 10^−5^*A dm*^−2^ computed from (18). *p*_*AT*_

*P* represents the thermodynamic limit of the system; in reality, it is unlikely to be achieved because the second law of thermodynamics implies that a fraction of the energy will be dissipated as heat.

#### The effect of the ionic conductances and extracellular impermeant molecules on energy consumption & p_min_

The leak conductance of the cell, in particular to the monovalent cations ions (*g*_*Na*_ and *g*_*K*_), exerts a strong effect on the energy consumption of the NKA. Figure 5 shows *p*_min_ as a function of *g*_*Na*_ and *g*_*K*_ when [*Na*^+^]_*e*_ = 147mM, [*K*^+^]_*e*_ = 3mM, *ν* = 3, and *κ* = 2. Also, we assume that *g*_*K*_ *> g*_*Na*_ since this is known to be so in most cells, as most cells have a resting potential less than -20mV [30]. To minimize *p*_min_, *g*_*Na*_ should be kept low. Note that when *g*_*Na*_ is small, *p*_min_ remains small for any value of *g*_*K*_. This means that when *g*_*Na*_ is small, *p*_min_ is less sensitive to any perturbations to *g*_*K*_.

**Fig. 5:**
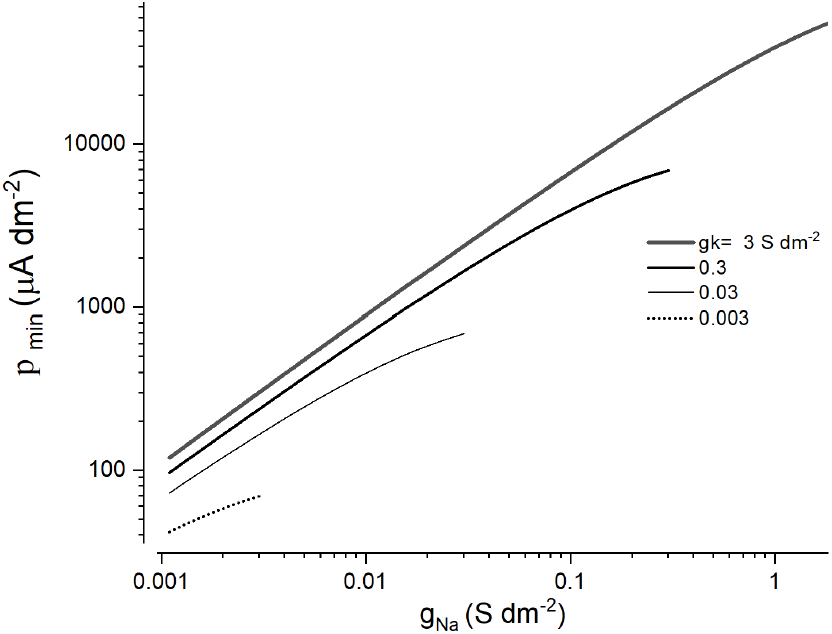
*p*_min_ as a function of *g*_*Na*_ is plotted for four different values of *g*_*K*_. *p*_min_ is calculated from (19). Note that we assume *g*_*Na*_ *< g*_*K*_, hence, each curve is plotted on a limited range for *g*_*Na*_.

## V. Active cells: Interaction of NKA and CCC transporters on cell regulation

In this section, we introduce three cation-coupled cotransporters into the pump-leak scenario (see Figure 1 for an illustration).

i. *K*^+^– *Cl*^−^ co-transporter (KCC), where one *K*^+^ and one *Cl*^−^ are transported out of the cell, with rate *c*_*KCC*_,
ii. *Na*^+^– *Cl*^−^ co-transporter (NCC), where one *Na*^+^ and one *Cl*^−^ are transported into the cell, with rate *c*_*NCC*_, and
iii. *Na*^+^– *K*^+^– *Cl*^−^ co-transporter (NKCC), where one *Na*^+^, one *K*^+^, and two *Cl*^−^ are transported into the cell with rate *c*_*NKCC*_.

The rate equations for the CCCs are obtained as follows (we only show it for KCC, the rest are similar to KCC). We assume a chemical equilibrium of the form [*K*^+^]_*i*_ + [*Cl*^−^]_*i*_ ⇔ [*K*^+^]_*e*_ + [*Cl*^−^]_*e*_ where the forward and backward rate constants are equal. Then:

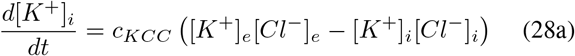

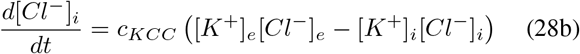

where *c*_*KCC*_ is a non-negative constant. Hence, in (6), the dynamics of each ion change as follows:

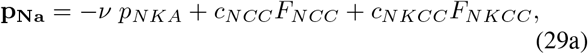

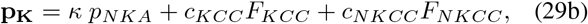

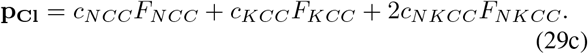

where *F*_*CCC*_ denotes the driving force of the CCC cotransporter and is described by:

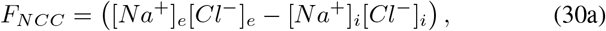

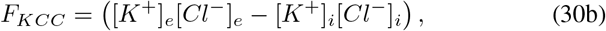

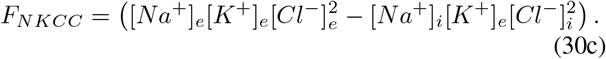

In this section, we examine the effect of adding all three types of CCC transporters separately to a cell with an active NKA transporter. All the CCCs rely on the ionic gradients established by the NKA to drive the system to a new steady state.

As we showed in Section III, in the absence of active transport, the cell moves into an equilibrium, where (as shown in (9))

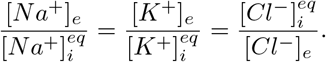

If a CCC is introduced into this passive cell, the current generated by the co-transporter becomes zero. We show this for the case of the KCC, the other co-transporters are similar. From the above equation 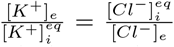 implies 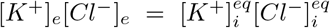, which (28) demonstrates that the ionic flux through the KCC co-transporter is zero. Similarly, all other CCC currents become zero. This implies that adding any of these co-transporters to the NKA-pump leak model with pump off can not change the equilibrium. However, they may change the rate at which the cell convergence to the steady state if the NKA is turned off [16].

In the classical PLM, since *Cl*^−^ is not actively transported, in the steady state, *E*_*Cl*_ is equal to the membrane potential, i.e. the transmembrane *Cl*^−^ gradient is at equilibrium. Hence *g*_*Cl*_ appears not to play much of a role in establishing the steady state (note it does not appear in the equations that determine the steady state). However, in the absence of a *Cl*^−^ conductance, the PLM is essentially frozen, since to preserve electroneutrality, [*Cl*^−^]_*i*_ needs to be able to change. It is also worth noting that in the PLM the magnitude of *g*_*Cl*_ can influence the rate at which the steady state is approached. When a CCC is introduced into the scenario, since *Cl*^−^ is now actively transported, the magnitude of *g*_*Cl*_ now becomes important.

In Figures 6-8, the steady state values of the ions concentrations, intracellular impermeant molecule, volume, and voltage are plotted at different co-transporter rates when only one co-transporter is active. To compute these steady states we solved a system of 5 algebraic equations, setting the right-hand side of (6a)-(6c) and (2) to zero and combining them with the constraint (4). The analytic expressions for these steady states are given in Appendix VIII-B

**Fig. 6:**
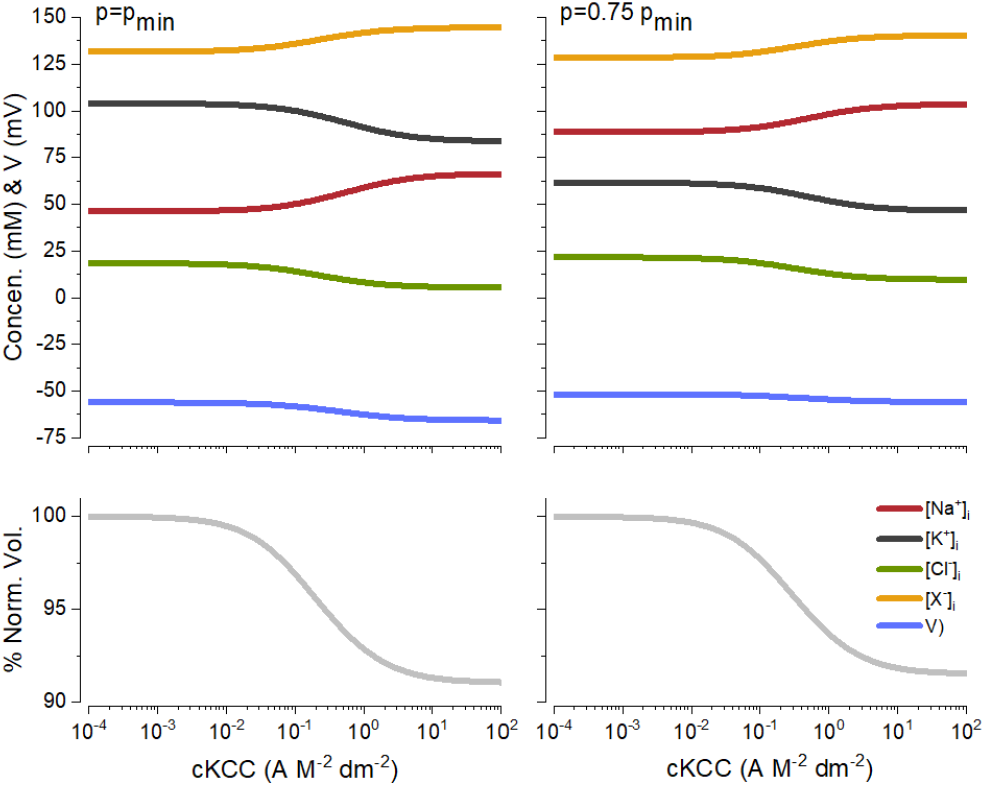
The steady state values of ion concentrations, intracellular impermeant molecule, volume, and voltage are plotted as a function of the KCC co-transporter rate. Other co-transporters are off and *g*_*Na*_ = 0.02*A*_0_, *g*_*K*_ = 0.03*A*_0_ and *g*_*Cl*_ = 0.002*A*_0_. (LHS) *p* = *p*_min_ and (RHS) *p* = 0.75*p*_min_.

Figures 6-8 are plotted for *p* = *p*_*min*_, the value of *p* where the cell takes its minimum volume and voltage at the steady state and for an intermediate value of *p, p* = 0.6*p*_min_ or *p* = 0.75*p*_min_. The behavior of the cell is qualitatively similar for other values of *p*. The results are not shown here.

In [15, Proposition 4.4 & Theorem 4.8], Mori proved that if (20) holds, for “sufficiently small” NKA and CCCs rates, namely *p*_*NKA*_ and *c*_CCC_, the PLEs with both active NKA and CCCs possess asymptotically stable steady states.^1^ Mori’s result does not guarantee uniqueness. Here, we derived these steady states in Appendix VIII-B, and the derivation guarantees the uniqueness. Since we have the explicit expressions for the steady states, similar to the case of PLEs with an active NKA in Section IV, we can find the range of *p*_*NKA*_ and *c*_CCC_ in which the steady states exist, i.e., we determine how small these parameters must be to ensure the existence of the steady states. Mori’s results ensure that our unique steady states are asymptotically stable for small *p*_*NKA*_ and *c*_CCC_. For larger *p*_*NKA*_ and *c*_CCC_, we show the stability numerically. A rigorous stability analysis of the steady states with arbitrary large *p*_*NKA*_ and *c*_CCC_ will be a topic for future investigations.

### K^+^– Cl^−^ co-transporter (KCC)

Figure 6 shows the influence of the KCC transporter on the steady state of a cell as the transport rate is increased at two different values of the NKA pump rate, *p*_min_ (left panels) and 0.75*p*_min_ (right panels). The other parameters are as in Table I. However, the sodium conductance has been increased accordingly since, at its default values, KCC exerts very little effect because both [*K*^+^]_*e*_ and [*Cl*^−^]_*e*_ are close to their equilibrium values. Note that *p*_min_ is computed for the new ion conductances based upon (19). KCC is an electroneutral transporter moving one *K*^+^ and one *Cl*^−^ in concert out of the cell, leading to a decrease of the concentration of both inside the cell, but it can only do so while adhering to the osmotic and charge constraints that govern the system. In what follows we explain how these constraints channel the behavior of the KCC, intuitively. An explicit formula of the steady states is given in Appendix VIII. Since for *z*_*X*_ = −1, the sum of (5) and (10) imply [*Na*^+^]_*e*_ +[*K*^+^]_*e*_ = 𝒪_*e*_/2, i.e, the sum of [*Na*^+^]_*i*_ and [*K*^+^]_*i*_ remain constant, [*Na*^+^]_*i*_ must increase as [*K*^+^]_*i*_ decreases (Figure 6, top panels). In addition, the intracellular electroneutrality ((5)) and osmotic balance ((10)) imply [*Cl*^−^]_*e*_ + [*X*]_*i*_ = 𝒪_*e*_/2.

Therefore, since the [*Cl*^−^]_*i*_ decreases, [*X*]_*i*_ must increase (Figure 6, top panels), while in turn, the volume decreases (Figure 6, bottom panels). To find out how *V* changes as *c*_*KCC*_ increases, we use (6a) at the steady state, i.e., when its right-hand side is zero:

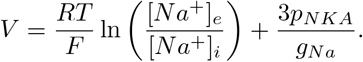

Since [*Na*^+^]_*i*_ increases as *c*_*KCC*_ increases, the voltage which is a decreasing function of [*Na*^+^]_*i*_, must decrease as *c*_*KCC*_ increases.

### Na^+^– Cl^−^ co-transporter (NCC)

It is first worth noting the commonalities of the CCC transporters; since they all use the *Na*^+^ and *K*^+^ gradient established by the NKA, their action always leads to an increase in [*Na*^+^]_*i*_ and a reciprocal decrease in [*K*^+^]_*i*_. In contrast to the KCC both the NCC and NKCC lead to increases in [*Cl*^−^]_*i*_ with a concomitant decrease in [*X*]_*i*_. In addition, both the NCC and NKCC lead to a depolarization of the membrane potential and an increase in cell volume, whereas opposites occur for the KCC. To maintain [*Na*^+^]_*e*_ + [*K*^+^]_*e*_ = 𝒪_*e*_/2, the *K*^+^ concentration must decrease, as shown in Figure 7 (top panels). From [*Cl*^−^]_*e*_ +[*X*]_*i*_ = 𝒪_*e*_/2 and the fact that [*Cl*^−^]_*i*_ increases, we conclude that [*X*]_*i*_ decreases and consequently the volume increases as the NCC transport rate increases, see Figure 7 (bottom panels). In this case, to analyze the voltage at the steady state we use (6b) and set its right hand to zero:

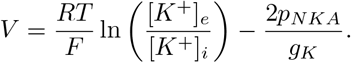

**Fig. 7:**
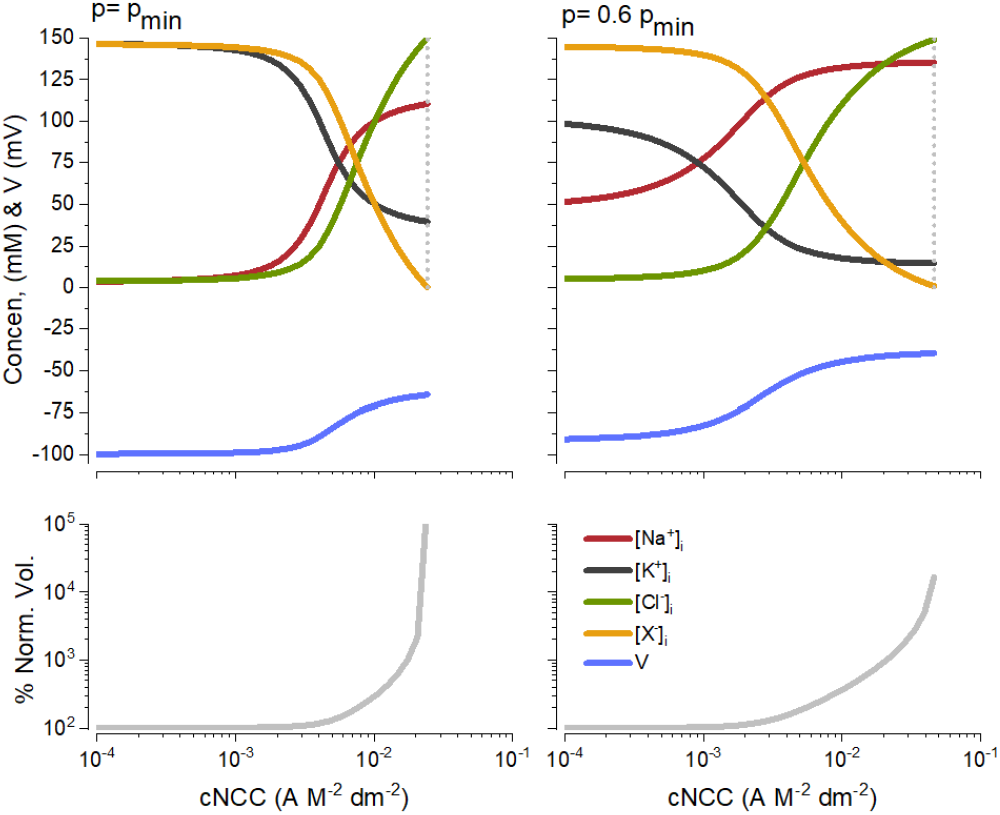
The steady state values of ion concentrations, intracellular impermeant molecule, volume, and voltage are plotted as a function of the NCC co-transporter rate. Other co-transporters are off. (LHS) *p* = *p*_min_ and (RHS) *p* = 0.6*p*_min_. The vertical dotted lines mark the value of *p* when the cell becomes unstable.

Since [*K*^+^]_*i*_ decreases and voltage is a decreasing function of [*K*^+^]_*i*_, voltage increases as [*K*^+^]_*i*_ decreases. Equivalently, voltage increases as *c*_*NCC*_ increases, see Figure 7 (middle panels). In contrast to the KCC, the action of the NCC can destabilize a cell. Above a critical value of *c*_*NCC*_ the volume does not stabilize, although the concentrations and voltage reach a steady state. As far as we know, this aspect of the NCC has not been noted before and only becomes evident when the NCC is modeled in the context where the membrane is water permeable and the membrane is pliant.

### Na^+^– K^+^– Cl^−^ co-transporter (NKCC)

As shown in Figure 8, the steady state behavior of a cell with an active NKCC is similar to a cell with an active NCC. When NKCC is active, one *Na*^+^, one *K*^+^, and two *Cl*^−^ enter the cell. So it is expected that the *Na*^+^ and *K*^+^ level increase, however, the sum of [*Na*^+^]_*i*_ and [*K*^+^]_*i*_ must remain constant. Hence, one of these ions must decrease. Since *g*_*K*_ is much higher than *g*_*Na*_, some *K*^+^ escape through the membrane, and therefore, the *K*^+^ level decreases. The rest of the characteristics of NKCC is similar to NCC cotransporter, except that it does not destabilize the cell volume.

**Fig. 8:**
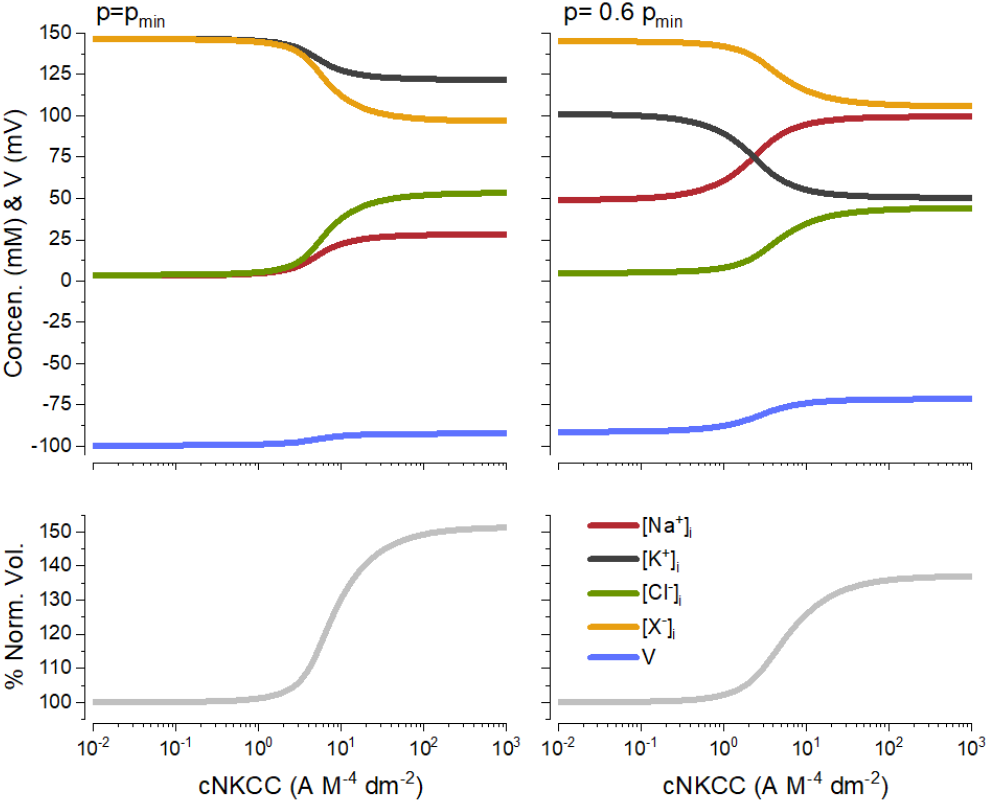
The steady state values of ion concentrations, intracellular impermeant molecule, volume, and voltage are plotted as a function of the NKCC co-transporter rate. Other co-transporters are off. (LHS) *p* = *p*_min_ and (RHS) *p* = 0.6*p*_min_.

The primary role of the CCCs appears to be to shift the [*Cl*^−^]_*i*_ away from equilibrium. In the case of KCC below equilibrium, while NCC and NKCC shift it above its equilibrium value. In addition, their activation can shift the steady state volume of the cell. In the case of KCC decreasing the volume, while in the case of NKCC and NCC increasing the volume. It is clear from our work that activation of either KCC or NKCC leads to predictable shifts in ion distributions without perturbing cellular stability. However, excessive activation of an NCC can lead to the loss of cellular stability. We have not investigated how the simultaneous deployment of different CCCs influences ionic distributions, leaving that for another time.

### Can CCC co-transporters be reversed?

The flux of ions through the co-transporters is determined by what we will term the “driving force”, *F*_*CCC*_*s* defined in (30). If the term is negative the ions are driven out of the cell (KCC), conversely, if the driving force is positive the ions are driven into the cell (NCC & NKCC). Notice that all of the co-transporters are symporters, i.e. all the transported ions always move together in the same direction.

We will use the analytical PLE to determine if it is possible to reverse the direction of the co-transporter [**?**], [31]. The concentrations within the driving force terms are primarily set by the NKA when the transport rate of the CCC is low. As the co-transporter rate is increased, the system is driven towards a steady state where the driving force term tends to zero. The steady state is determined by the competition between the NKA and the CCCs.

In what follows we show that at the steady state, the driving forces generated by the CCC co-transporters do not change signs. First, we show that the steady state value of the KCC driving force is always negative. For *c*_*KCC*_ = 0, the steady state value of the driving force is calculated from (17a) and (17c) as follows.

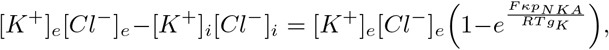

that is always negative since 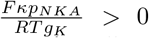 for positive parameters, hence driving ions out of the cell. For *c*_*KCC*_ > 0, using (38b) and (38c) given in Appendix VIII-B, the driving force becomes 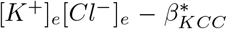, where 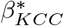 is the unique solution of (32) and can be computed numerically.

Figure 9 (top panel), shows the KCC driving force as a function of *c*_*KCC*_ for three different values of the NKA pump rate. As *c*_*KCC*_ increases, the driving force remains negative and never reverses.

**Fig. 9:**
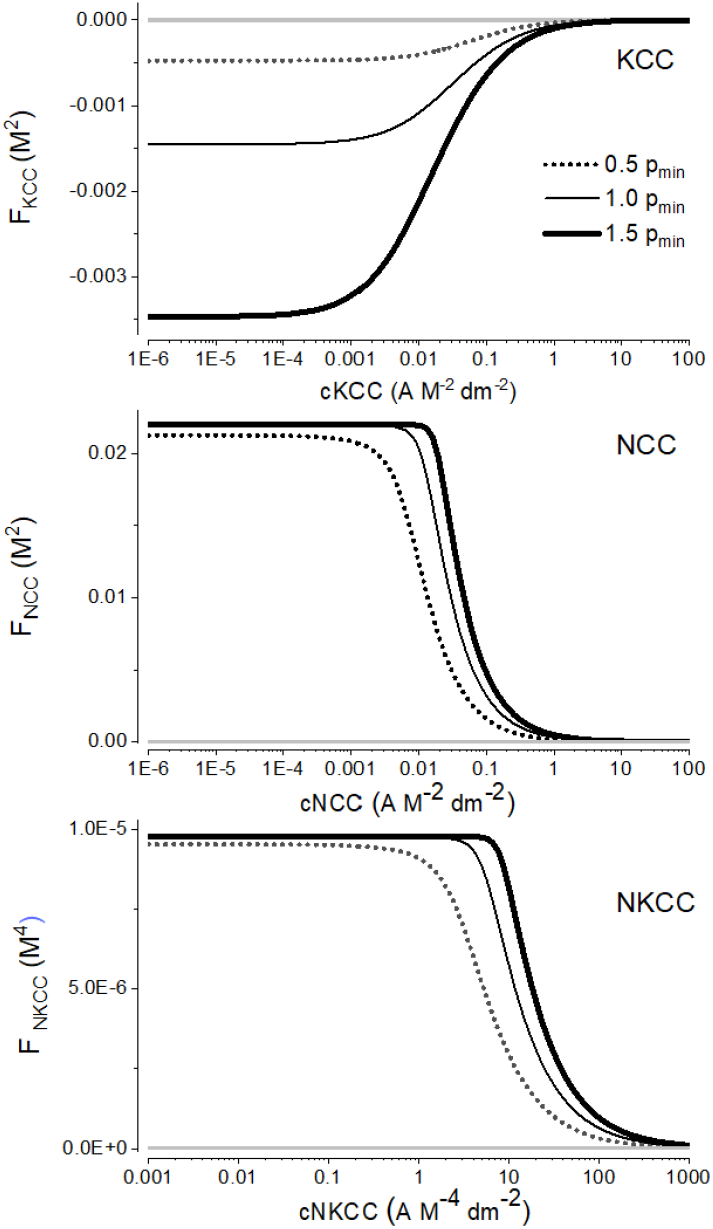
The steady state values of CCC driving forces as functions of *c*_*CCC*_ are shown for three values of *p, p* = *p*_min_, 0.5*p*_min_, 1.5*p*_min_. KCC’s driving force is always negative, while NCC and NKCC are always positive.

Then, we show that the steady state value of the NCC driving force is always positive. For *c*_*NCC*_ = 0, the steady state value of the driving force is calculated from (17a) and (17b):

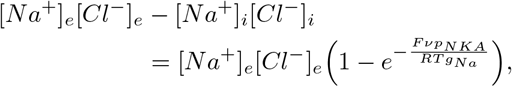

that is always positive since 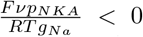 for positive parameters. For *c*_*NCC*_ > 0, using (38a) and (38c) given in Appendix VIII-B, the driving force becomes 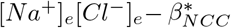, where 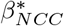 is the unique solution of (34) and can be computed numerically. Figure 9 (middle panel) depicts the NCC driving force 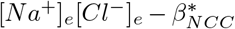 as a function of *c*_*NCC*_. As *c*_*NCC*_ increases, although the force decreases, it preserves its sign and remains positive.

Finally, we show that the steady state value of the NKCC driving force is positive if 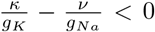 and is negative, otherwise. For *c*_*NKCC*_ = 0, the steady state value of the driving force can be calculated from (17a)–(17c):

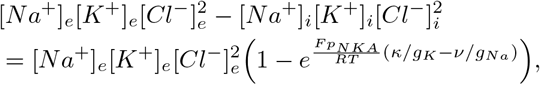

that is positive if 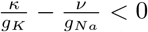 and is negative, otherwise. For the given parameters in Table I, this value is always negative. For *c*_*NKCC*_ > 0, using (38a)–(38c) given in Appendix VIII-B, the driving force becomes 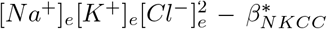, where 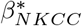 is the unique solution of (36) and can be computed numerically. Figure 9 (lower panel) depicts the NKCC driving force 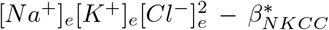 increases, although the force decreases, it always remains positive and does not change signs.

It is worth noting that all of the CCCs preserve their default direction of ion transport, regardless of the choice of the parameters for the PLE, so long as the parameters generate a stable system. The CCCs could be reversed if the ion concentrations are perturbed away from their steady state values, but the system will return to a steady state with the CCCs transporting ions in their default direction.

## VI. Discussion

Instead of attempting to model a particular kind of cell, we have explored what sorts of steady state ion distributions can be achieved by combing the classical PLEs with CCCs and extracellular impermeant molecules. We have also identified some mathematical principles that undergird such systems.

There are several analytical tools for making sense of how voltage-gated ion channels interact to generate neuronal excitability [32], [33], but little for the case where one incorporates ion transporters. The analysis becomes a lot more complicated as soon as one allows for the accumulation of ions and the flux of water driven by changes in osmolarity.

Analytical solutions to models have a significant advantage over piecemeal simulation since they can make evident the dependence of the system’s behavior on its parameters. Additionally, in simulations of a system, one is in a sense working blind, while analytical solutions give one an overview of the system.

Whether the steady state of a model of cellular ion transport is stable or not is a matter of great importance in evaluating models. In [34], Weinstein studied “local” stability by linearization about a physiologic reference condition. The framework introduced by Mori of viewing PLEs as being governed by a free energy principle is very useful. It allows one to get a global overview of the system and predict what behaviors may or may not occur without computing the steady states. However, it cannot fully characterize the full range of parameters in which the steady states exist and are stable. In this work, we were able to derive the steady states explicitly and found the full range of the parameters in which the steady states exist and numerically showed that these steady states are stable. Mori’s work ensures “global” stability of the steady states for PLEs with a constant pump, however, it cannot determine their global stability in the presence of co-transporters. In this work, we explicitly derived unique steady states and numerically showed their stability. A rigorous stability analysis will be left for another time. In [35], [36], the authors go beyond stability analysis and apply linear optimal control theory to stabilize *Na*^+^ flux, cell volume, and cell pH.

### The effect of NKA stoichiometry on energy utilization

The NKA utilizes approximately 20-30 percent of all cells’ energy budget, its continuous action is required to forestall an osmotic catastrophe induced by the presence of impermeant molecules. This is an ongoing process that does not relent.Given that the NKA constitutes so much of a cell’s energy budget, cells have probably evolved mechanisms to constrain energy consumption. If the pump rate is too low small fluctuations in the pump rate will lead to large ones in cell volume. See Figure 3 lower right panel.

We have shown that if one incorporates the thermodynamics of ion transport into the PLEs, a natural limit to the rate of ATP consumption emerges which prevents excessive ATP consumption, see (27) and Figure 4. This to the best of our knowledge has not been demonstrated before and it is a feature that only becomes apparent if the action of the NKA is viewed in the framework of the PLEs in conjunction with the energy of the pump.

A complete model of the NKA would require a characterization of all of the steps in its cycle, with their dependence on ion concentration and voltage. However, we have shown in Section VIII that if one is only interested in the steady state of the PLEs, the precise form of the NKA is of no importance. Where it becomes important is in the system’s transient response, which we have not considered here.

Theoretically, we can contemplate any stoichiometry, however, the constraints of protein chemistry probably limit the number of ions that can be coordinated at one time. So, for example, there are no cases where more than five ions or molecules are transported during one cycle of a transporter.

### Water permeability

Many cellular transport models omit a crucial aspect (with some exceptions), namely water movement. It is certainly possible to simulate the system without this but it runs the risk of arriving at a model which might be unstable if water transport is included, as it ultimately must since all membranes are water permeable.

The omission of water permeability may veil an instability in the system. One may find that a model produces the correct electrical phenomenology. However, the volume may be unstable when one introduces water permeability and cell pliancy.

### The effect of leak conductances on energy consumption and cellular stability

The values of *g*_*K*_ and *g*_*Na*_ are primary determinants of ATP utilization. Moreover, although this is not well known, leak conductances can strongly impact the action potential threshold [37]. But little is known about how cells set and regulate their resting input conductances. If the input conductance is very low, the opening of ion channels can lead to pronounced changes in voltage which may be damaging.

Thus, cells need to keep their input resistance relatively low to avoid large swings in membrane potential, which could rupture the membrane. Since ion channels open in a stochastic fashion, varying ion fluxes are inevitable. If the input resistance is too high, the transport of metabolites like electrogenic co-transporters will induce significant changes in the membrane potential [38].

There is a large class of channels that are open at rest that determines the baseline permeability of cells and their input resistance. These include, *K*_2*P*_ channels [39], NALCN channels [40], Kir channels [41], and HCN channels [42].

### Extracellular impermeantions

In this publication, we have introduced a simple extension to the PLEs that allows the development of a stable potential and volume by introducing impermeant extracellular molecules. If the NKA is inactive, the system represents a stable Donnan equilibrium at thermodynamic equilibrium. This stratagem of using extracellular molecules to stabilize the cell has only been mentioned by Fraser and Huang [8]. Although impermeant extracellular molecules could stabilize a cell, it does not account for the asymmetric *Na*^+^ and *K*^+^ distribution encountered in actual cells. Moreover, it would require a very high concentration of *Y* (50 − 100 mM) to sustain a comparable concentration of *X*, which is needed for normal cell function.

Many years ago, Donnan [43], among others, thought that cells might be in a passive Donnan equilibrium. However, he did not consider the possible influence of impermeant extracellular molecules. Subsequently, several investigators demonstrated that the operation of the NKA sustains a stable steady volume through the PLEs. We are unaware of any cases where cells have been shown to be in a stable Donnan equilibrium sustained by a high concentration of impermeant extracellular molecules. However, some organisms may use this strategy when aestivating [44], where energy utilization must be kept very low.

### CCCs

The addition of the CCCs cannot change the osmotic and electroneutrality constraints. Moreover, the CCCs do not change the qualitative dynamics of the system. We showed that if a CCC is added to stable PLEs, it cannot induce qualitatively different dynamics, i.e., PLEs remain to possess unique globally stable steady states. However, it is possible that the addition of a CCC could change the rate at which the cell converges to a new steady state.

A novel finding that emerged from our study is that NCC can induce cellular instability at high pump rates. The overall effects of the NKCC and NCC are the same; increasing [*Na*^+^]_*i*_, [*Cl*^−^]_*i*_, and volume. Therefore it is something of a puzzle why there are two mechanisms. It is interesting to note that NCC has a very restricted expression, only being found in the kidney and bone. Although NKCC2 is only found in the kidney, NKCC1 has widespread expression [4], [45]. NKCC can be considered to have what might be thought of as a safety mechanism that limits the level of [*Cl*^−^]_*i*_ that can be achieved and the cell volume expanding uncontrollably. This case of the NCC illustrates the power of theory, which allows one to go beyond intuitions to make explicit and unexpected predictions.

Neuroscientists often say that the role of the NKA is to generate the resting potential, which provides the foundation for generating cellular excitability. However, the NKA’s primary role appears to oppose the Donnan effect to achieve cellular stability. It is worth bearing this in mind when considering other transporters’ roles since their roles may be rather subtle.

It is by no means obvious that the incorporation of CCCs into the PLM would not lead to the development of novel dynamics. In this paper, we have shown that they cannot do so, and that the parameter space of this system does not harbor regions with exotic dynamics. An analytical approach makes evident features of the system that would not be apparent in the absence of explicit mathematical expressions. This allowed us to show that the CCCs are unlikely to reverse and reveal the precise coupling between the NKA and ATP utilization, as well as the number of new features of the PLM.

## VII. Materials and Methods

All computations were performed using MATLAB, release R2020b (The MathWorks, Inc., Natick, MA). All graphs were produced with OriginPro (OriginLab, Northampton, MA).

A small capacitance (*C*) allows rapid changes in voltage. In this case, the PLEs become “stiff,” and special numerical solvers are required to solve this system. In this work, for the case of the non-linear NKA models, the system of equations was solved using the MATLAB stiff differential equation solver ode15s or ode23tb (the results are similar, only their speeds are different for different parameters.)

## VIII. Appendix

### A. Various mathematical models for NKA pump mechanisms

In this section, we show that although the stoichiometry of the pump effect the steady states, the precise mathematical models of NKA do not change them qualitatively. In addition, the precise mathematical models of NKA have no impact on ATP consumption. To this end, we consider a constant model and a nonlinear model as follows.

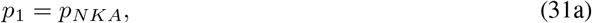

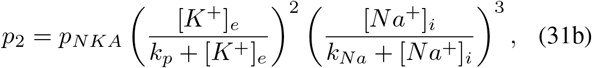

where *p*_*NKA*_ = *pA*_0_ is a constant. *k*_*p*_ and *k*_*Na*_ are apparent dissociation constants for *Na*^+^ and *K*^+^ and are equal to *k*_*p*_ = 0.883 mM and *k*_*Na*_ = 3.56 mM.

In Figure 10 we compare the ATP consumption rate of the constant and nonlinear pumps, namely, 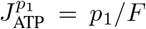 and 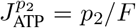 as a function of the pump rate *p*. Notice that for the latter, the ATP consumption rate comes close to an asymptotic value when *p* ≈ 0.75 *n mol*^−1^ *dm*^−2^, but carries on increasing very slowly.

**Fig. 10:**
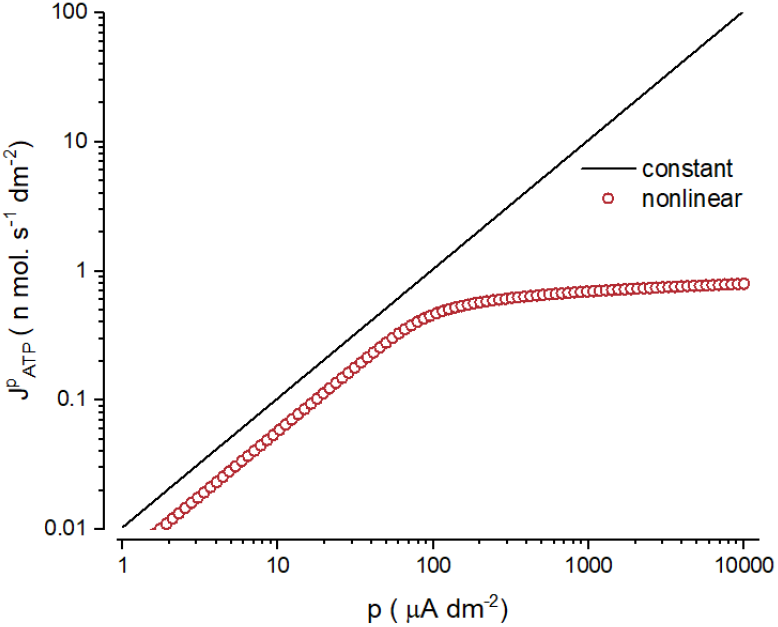
The ATP consumption rate as a function of the pump rate, for the constant NKA model given in (31a) and the nonlinear NKA model given in (31b).

In Figure 11 we plot the steady state of the system as a function of the ATP consumption rate for the constant and the nonlinear models of NKA. As the figure depicts, the steady state values coincide exactly. We observed that this also occurs for less complicated nonlinear dynamics such as *p*_*NKA*_[*Na*^+^]_*i*_ [16], *p*_*NKA*_ ([*Na*^+^]_*i*_/[*Na*^+^]_*e*_)^3^ [14], and *p*_*NKA*_ ([*K*^+^]_*e*_/[*K*^+^]_*i*_)^2^ ([*Na*^+^]_*i*_/[*Na*^+^]_*e*_)^3^ [46] (the result is not shown here). Therefore, the precise form of the NKA model is not important in determining the steady state of PLEs.

**Fig. 11:**
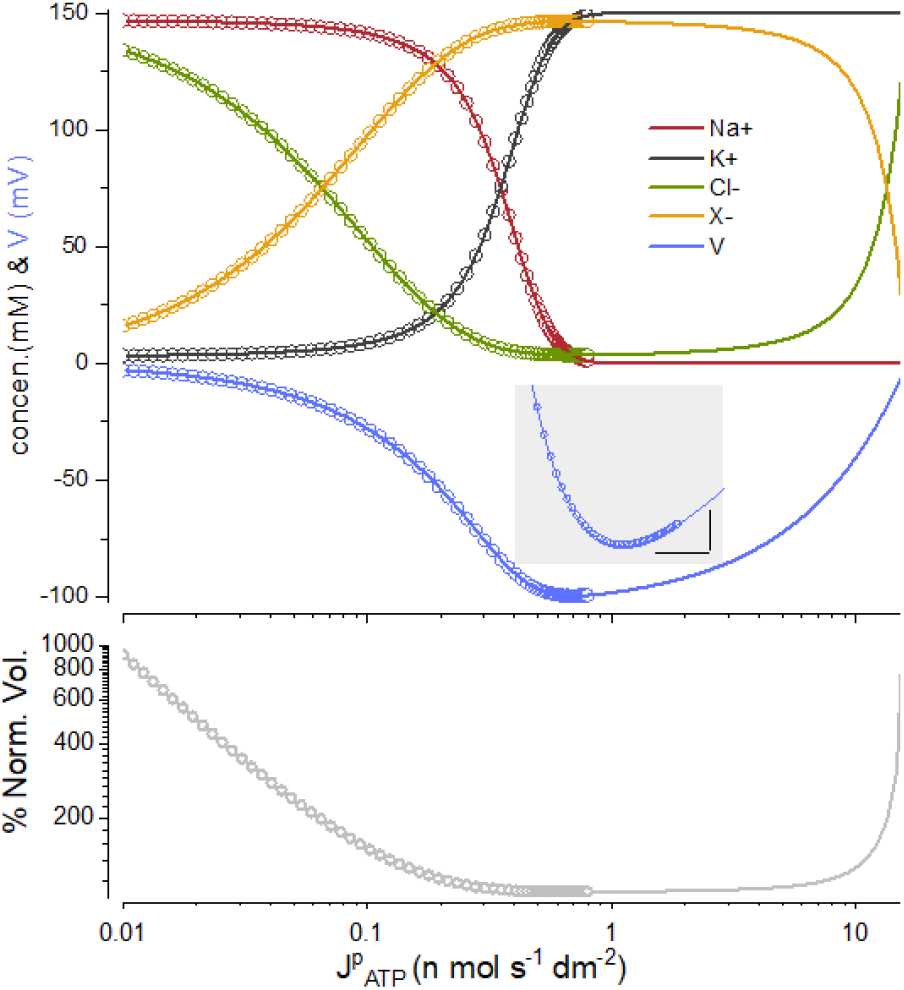
Steady state values as a function of the ATP consumption rate, for the constant NKA given in (31a) (solid curve) and the nonlinear NKA model given in (31b) (circles). The inset shows the nadir of the voltage with scale bars of 0.5*mV* and 0.1 *n mol s*^−1^*dm*^−2^.

### B./ Steady state values of PLEs in the presence of one co-transporter

In this section, we derive the steady state values of a PLM in the presence of a constant NKA pump and one CCC cotransporter. To compute the steady states, we let the righthand side of (6a)-(6c) and (2) to zero with constraint (4). The analytic expressions of these steady states are as follows.

For the PLEs with an active KCC, let

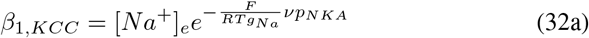

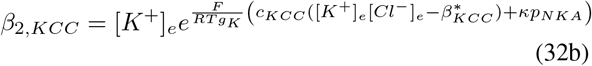

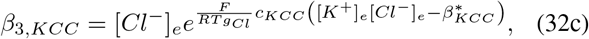

where 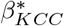 is the unique solution, *x*, which can be calculated numerically from the following equation:

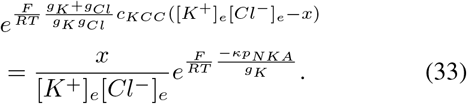

For the PLEs with an active NCC, let

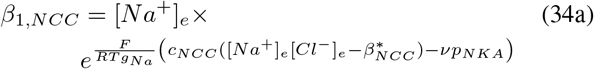

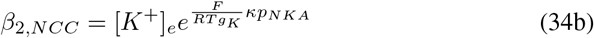

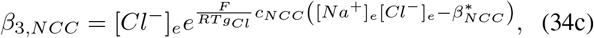

where 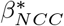 is the unique solution of

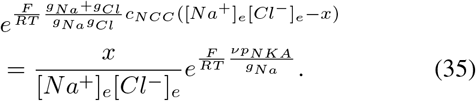

For the PLEs with an active NKCC, let

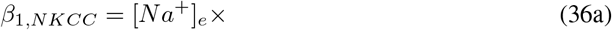

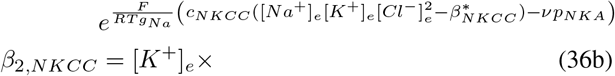

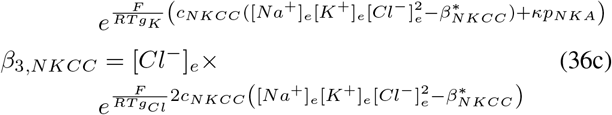

where 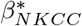 is the unique solution of

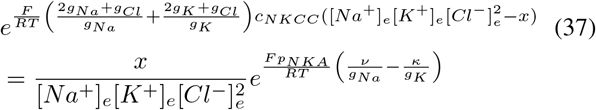

Note that since the left-hand side of (33) is a strictly decreasing function of *x* and its right-hand side is a strictly increasing and linear function of *x*, the intersection must be unique. For the given range of parameters, the right and lefthand sides intersect at exactly one point which determines the existence of exactly one steady state of PLEs. For the same reason, PLEs with an active NCC or NKCC possess a unique steady state, as derived below.

The unique steady state values of the PLEs with one CCC co-transporter are

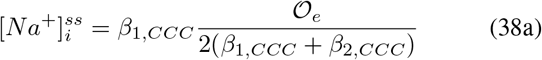

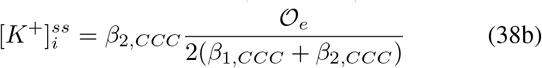

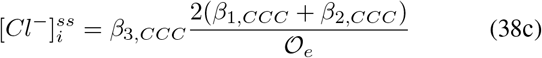

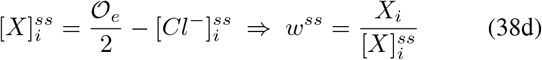

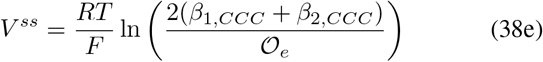

where *β*_*i,CCC*_ is either *β*_*i,KCC*_, *β*_*i,NCC*_, or *β*_*i,NKCC*_.

Similar to the case with only an active NKA pump, to have a steady state with positive volume, 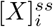 must be positive. Simple calculations can give a range of parameters *p* and *c*_*CCC*_ that make 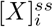 positive. Here, for *p* = *p*_min_, we observed that 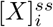 is positive for any values of *c*_*KCC*_ and *c*_*NKCC*_. However, for *c*_*NCC*_ there is a value that 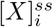 becomes zero and then negative. This value is shown by the vertical dotted line in Figure 7. In summary, unlike in [15] which the existence of the steady states is proved for a small amount of *p* and *c*_*CCC*_, we show the existence of the steady states for any possible values of *p* and *c*_*CCC*_.

These steady states are plotted in Figures 6-8 as *c*_*CCC*_ varies and *p* = *p*_*min*_ and *p* = 0.6*p*_*min*_. All other parameters are fixed. First we numerically solve (33), (35), and (37) for 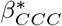. Then plugging 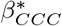 into (32), (34), (36), respectively, to compute 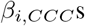 and finally, using these 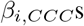 we compute the steady states values given in (38).

## Acknowledgment

We dedicate this paper to the memory of Alexey Verninov. We thank T. Budde for helpful advice. Z.A. was supported by Simon Foundations 712522 and Z.A. and A.R.K. were supported by NSF 2037828.

## Footnotes

1 Indeed, in [15, Proposition 4.4 & Theorem 4.8], these results are shown for a nonlinear active pump which does not depend on the intracellular concentrations. However, these results can easily be generalized to cotransporters that depend on the intracellular concentrations.

## References

[1] D C Tosteson and J F Hoffman. Regulation of cell volume by active cation transport in high and low potassium sheep red cells. J Gen Physiol, 44:169–94, 1960.

[2] F.G. Donnan. Theorie der membrangleichgewichte und membranpotentiale bei vorhandensein von nicht dialysierenden elektrolyten. ein beitrag zur physikalisch-chemischen physiologie. Zeitschrift ftir Elektrochemie und angewandte physikalische Chemie, 17:572–581, 1911.

[3] Juan Pablo Arroyo, Kristopher T Kahle, and Gerardo Gamba. The slc12 family of electroneutral cation-coupled chloride cotransporters. Molecular Aspects of Medicine, 34(2-3):288–298, 2013.

[4] E. Delpire and K. B. Gagnon. Na^+^− K^+^− 2Cl^−^ cotransporter (NKCC) physiological function in nonpolarized cells and transporting epithelia. Compr Physiol, 8(2):871–901, 2018.

[5] Thomas A. Chew, Benjamin J. Orlando, Jinru Zhang, Naomi R. Latorraca, Amy Wang, Scott A. Hollingsworth, Dong-Hua Chen, Ron O. Dror, Maofu Liao, and Liang Feng. Structure and mechanism of the cation–chloride cotransporter nkcc1. Nature, 572(7770):488–492, 2019.

[6] E Jakobsson. Interactions of cell volume, membrane potential, and membrane transport parameters. Am J Physiol, 238(5):C196–206, 1980.

[7] Julio A Hernández. A general model for the dynamics of the cell volume. Bull Math Biol, 69(5):1631–48, 2007.

[8] J.A. Fraser and L.-H. Huang C. Quantitative techniques for steady-state calculation and dynamic integrated modelling of membrane potential and intracellular ion concentrations. Prog. Biophys. Mol. Biol., 94(3):336–72, 2007.

[9] K. Dijkstra, J. Hofmeijer, S. A. van Gils, and M. J. van Putten. A biophysical model for cytotoxic cell swelling. J Neurosci, 36(47):11881–11890, 2016.

[10] I. A. Vereninov, V. E. Yurinskaya, M. A. Model, F. Lang, and A. A. Vereninov. Computation of pump-leak flux balance in animal cells. Cell Physiol Biochem, 34(5):1812–23, 2014.

[11] V. L. Lew, H. G. Ferreira, and T. Moura. The behaviour of transporting epithelial cells. i. computer analysis of a basic model. Proc R Soc Lond B Biol Sci, 206(1162):53–83, 1979.

[12] V. E. Yurinskaya, A. A. Rubashkin, and A. A. Vereninov. Balance of unidirectional monovalent ion fluxes in cells undergoing apoptosis: why does na plus /k plus pump suppression not cause cell swelling? Journal of Physiology-London, 589(9):2197–2211, 2011.

[13] K. Sharp, E. Crampin, and J. Sneyd. A spatial model of fluid recycling in the airways of the lung. J Theor Biol, 382:198–215, 2015.

[14] J. Keener and J Sneyd. Mathematical Physiology I: Cellular Physiology. Springer, New York, NY, 2nd. edition, 2009.

[15] Y. Mori. Mathematical properties of pump-leak models of cell volume control and electrolyte balance. Journal of Mathematical Biology, 65(5):875–918, 2012.

[16] V. E. Yurinskaya, I. A. Vereninov, and A. A. Vereninov. Balance of Na^+^, K^+^, and Cl^−^ unidirectional fluxes in normal and apoptotic u937 cells computed with all main types of cotransporters. Front Cell Dev Biol, 8:591872, 2020.

[17] R. P. Garay and P. J. Garrahan. The interaction of sodium and potassium with the sodium pump in red cells. The Journal of Physiology, 231(2):297–325, 1973.

[18] N P Smith and E J Crampin. Development of models of active ion transport for whole-cell modelling: cardiac sodium-potassium pump as a case study. Progress in Biophysics and Molecular Biology, 85(2-3):387–405, 2004.

[19] D. F. Rolfe and G. C. Brown. Cellular energy utilization and molecular origin of standard metabolic rate in mammals. Physiol Rev, 77(3):731–58, 1997.

[20] Yizeng Li, Xiaohan Zhou, and Sean X Sun. Hydrogen, bicarbonate, and their associated exchangers in cell volume regulation. Frontiers in Cell and Developmental Biology, 9:1640, 2021.

[21] A.M. Weinstein and J.L. Stephenson. Electrolyte transport across a simple epithelium. steady-state and transient analysis. Biophysical Journal, 27(2):165–186, 1979.

[22] Mariia Dvoriashyna, Alexander J.E. Foss, Eamonn A. Gaffney, Oliver E. Jensen, and Rodolfo Repetto. Osmotic and electroosmotic fluid transport across the retinal pigment epithelium: A mathematical model. Journal of Theoretical Biology, 456:233–248, 2018.

[23] W.F. Boron and E.L. Boulpaep. Medical Physiology: A cellular and molecular approach. Elsevier, 3rd edition, 2016.

[24] A. Varghese and G. R. Sell. A conservation principle and its effect on the formulation of Na-Ca exchanger current in cardiac cells. J Theor Biol, 189(1):33–40, 1997.

[25] A. R. Kay. How cells can control their size by pumping ions. Frontiers in Cell and Developmental Biology, 5(41), 2017.

[26] R F Burton. The composition of animal cells: solutes contributing to osmotic pressure and charge balance. Comp Biochem Physiol, B, 76(4):663–71, 1983.

[27] Peter Läuger. Electrogenic ion pumps. 1991.

[28] R. Milo and R. Phillips. Cell Biology by the Numbers. Garland Science, New York, NY, 2015.

[29] Ronald J. Clarke, Michelina Catauro, Helge H. Rasmussen, and Hans-Jürgen Apell. Quantitative calculation of the role of the na+,k+-atpase in thermogenesis. Biochimica et Biophysica Acta (BBA) - Bioenergetics, 1827(10):1205–1212, 2013. OA status: bronze.

[30] T.F. Weiss. Cellular Biophysics: Transport, volume 1. MIT Press, Boston, MA, 1996.

[31] Yasuhiro Kakazu, Soko Uchida, Takashi Nakagawa, Norio Akaike, and Junichi Nabekura. Reversibility and cation selectivity of the k-cl cotransport in rat central neurons. Journal of Neurophysiology, 84(1):281–288, 2000.

[32] D. Sterrat, B. Graham, A. Gillies, and D. Willshaw. Principles of Computational Modelling in Neuroscience. Cambridge University Press, 2011.

[33] Eugene M Izhikevich. Dynamical systems in neuroscience. MIT press, 2007.

[34] A.M. Weinstein and E. D. Sontag. Dynamics of cellular homeostasis: Recovery time for a perturbation from equilibrium. Bulletin of Mathematical Biology, 59(3):451–481, 1997.

[35] A.M. Weinstein and E. D. Sontag. Modeling epithelial cell homeostasis: Assessing recovery and control mechanisms. Bulletin of Mathematical Biology, 66(5):1201–1240, 2004.

[36] A.M. Weinstein and E. D. Sontag. Modeling proximal tubule cell homeostasis: Tracking changes in luminal flow. Bulletin of Mathematical Biology, 71(6):1285–1322, 2009.

[37] A. R. Kay. What gets a cell excited? kinky curves. Adv Physiol Educ, 38(4):376–80, 2014.

[38] N. Berndt and H. G. Holzhutter. The high energy demand of neuronal cells caused by passive leak currents is not a waste of energy. Cell Biochem Biophys, 67(2):527–35, 2013. Berndt, Nikolaus Holzhutter, Hermann-Georg eng 2013/03/13 06:00 Cell Biochem Biophys. 2013 Nov;67(2):527–35. doi: 10.1007/s12013-013-9538-3.

[39] Péter Enyedi and Gábor Czirják. Molecular background of leak k+ currents: two-pore domain potassium channels. Physiological reviews, 90(2):559–605, 2010.

[40] D. Ren. Sodium leak channels in neuronal excitability and rhythmic behaviors. Neuron, 72(6):899–911, 2011.

[41] CG Nichols and AN Lopatin. Inward rectifier potassium channels. Annual review of physiology, 59(1):171–191, 1997.

[42] C. Wahl-Schott and M. Biel. Hcn channels: Structure, cellular regulation and physiological function. Cellular and Molecular Life Sciences, 66(3):470–494, 2009.

[43] FG Donnan. Linkage of physico-chemical processes in biological systems. Nature, 149(3779):383, 1942.

[44] Kenneth B. Storey and Janet M. Storey. Molecular physiology of freeze tolerance in vertebrates. Physiological Reviews, 97(2):623–665, 2017.

[45] Arohan R. Subramanya. Thiazide-Sensitive NaCl Cotransporter, pages 57–92. Springer International Publishing, 2020.

[46] Santosh Manicka and Michael Levin. Modeling somatic computation with non-neural bioelectric networks. Scientific Reports, 9(18612), 2019.

